# A selective force driving metabolic genes clustering

**DOI:** 10.1101/2022.09.05.506644

**Authors:** Marco Fondi, Francesco Pini, Christopher Riccardi, Pietro Gemo, Matteo Brilli

**Author notes:** **Corresponding author:** E-mail: MB -, MF -, FP.

## Abstract

The evolution of operons has puzzled evolutionary biologists since their discovery and many theories exist to explain their emergence and spreading. The presence of several plausible hypotheses dealing with operon emergence/evolution/spreading is indicative of the absence of a universal causal factor for this evolutionary process. Here, we argue that the way in which DNA replication and cell division are coupled in microbial species introduces an additional selective force that may be responsible for the clustering of functionally related genes on chromosomes. We interpret this as a preliminary and necessary step in operon formation. Specifically, we start from the observation that during DNA replication differences in copy number of genes that are found at distant loci on the same chromosome arm exist. We provide theoretical considerations suggesting that, when genes of the same metabolic process are far away on the chromosome, this results in perturbations to metabolic homeostasis. By formalizing the effect of DNA replication on metabolic homeostasis based on Metabolic Control Analysis, we show that the above situation provides a selective force that can drive the formation of gene clusters and operons. Finally, we confirmed that, in present-day genomes, this force is significantly stronger in those species where the average number of active replication forks is larger and quantify the theoretical contribution of this feature on the distribution of extant gene clusters and operons.

## Introduction

Operons (Jacob and Monod, 1961) are one of the hallmarks of prokaryotic genomes; in their most basic form they comprise a single promoter at the 5′ end, followed by two or more genes and a transcription terminator at the 3′ end, therefore they are transcribed into polycistronic messenger RNAs. Additional regulatory sites (alternative and/or internal promoters, attenuators etc.) can be present to provide fine control over the gene expression levels (Lawrence, 2003; Zheng *et al*., 2002); genes in an operon often participate to a same functional process. While common in Prokaryotes, they are exceptions (Blumenthal *et al*., 2002; Boycheva *et al*., 2014; Lewis, 1978) or peculiarities (Bussotti *et al*., 2021) in Eukaryotes, a fact that was explained by the smaller effective population sizes of many eukaryotes that - according to the *drift model*, would lead to operon disruption (Lynch, 2006).

The evolution of operons is debated since their discovery; early ideas often coupled operon formation and metabolic pathway evolution but, while metabolic pathways are often ancient, taxonomic variability in operon organization of the corresponding genes suggests a more recent evolutionary history (e.g. Fani *et al*. (2005)).

The process of operon evolution can be split into two aspects: one is *operon assembly*, by which scattered genes form clusters that may then evolve into operons. The second is *operon maintenance* in evolution, which depends on the fitness differential provided by the operon with respect to the same genome with scattered genes. Since what makes an operon advantageous *once* it is formed is not necessarily what drove the genes at nearby loci, *assembly* and *maintenance* may be consequences of very different forces. For instance, an operon may reduce noise in protein abundances as previously suggested (Ray and Igoshin, 2012), but this advantage can probably be efficiently selected for by evolution only *after* the operon has formed (the same authors refer to *operon maintenance*) and to a much lesser extent during the intermediate steps. Models dealing with the identification of the force driving gene clustering during the formation of operons, exist. The Fisher model suggests that the compactness of operons may reduce the chances of destructive recombination events (Stahl and Murray, 1966) an idea recently revived in (Fang *et al*., 2008) where the authors, using simulations, also showed that random gene deletion could drive the formation of gene clusters. The co-regulation model is intrinsically linked with the operon rationale: genes stay together to facilitate their coordinated expression (Price *et al*., 2005), but others have shown that the formation of operons for the purpose of co-regulation is both unnecessary and implausible (Lawrence and Ochman, 1998; Lawrence and Roth, 1996) because independent promoters can evolve characteristics enabling co-expression of genes encoded at different loci. The *selfish operon model* (Lawrence and Roth, 1996) focuses on the formation of operons by deletion of intervening genes after horizontal gene transfer of a much larger fragment. However, this hypothesis was criticized on the basis of a high degree of gene clustering for housekeeping genes, whose horizontal transfer is infrequent (Pál and Hurst, 2004). Another possible explanation for the evolution of operons is that the coupling of transcription and translation in Prokaryotes, together with a strong limitation of diffusion by crowding, would confine products in a relatively small volume of the cytoplasm, maximizing interactions and metabolic fluxes (Dandekar *et al*., 1998; Shieh *et al*., 2015; Sneppen *et al*., 2010). A similar idea forms the basis for the Protein Immobility Model (Svetic *et al*., 2004) where gene clustering is beneficial because it reduces metabolic costs. The main assumption is that large enzymes don’t diffuse appreciably in the cell i.e. in this model metabolic sources and sinks are at fixed positions in the cell. Under this view, a gradient of the intermediate builds up which is proportional to the squared distance separating source and sink. It is the increased amount of metabolites required to sustain the flux through the pathway at increasing enzyme distances that provides the selective pressure for the formation of gene clusters under the PIM. However, not all proteins encoded by the same operon interact, channeling of metabolic intermediates is not so widespread, and recent estimates, withstanding the reduced mobility of proteins in the cell with respect to pure water, suggest that two particles can find each other in the cell in a matter of seconds (Schavemaker *et al*., 2018), a much shorter time scale than the characteristic time of metabolic systems. By using stochastic simulations of simple biochemical systems, evidences were found for noise reduction in the abundance of proteins encoded by operons, however this was limited to some type of interaction established by the products (Ray and Igoshin, 2012). It should be added that the two hypotheses mentioned above strongly rely on the coupling of transcription and translation that might not be as general as previously assumed (Irastortza-Olaziregi and Amster-Choder, 2021). Genome size and/or its proxies were also suggested to be markers of the intensity of natural selection for operon organization, under the idea that since larger genomes have more complex genetic networks thanks to the presence of more transcription factors, they would endure weaker selection for operons than smaller genomes, where regulation alternatives are scarce (Nuñez *et al*., 2013).

Several hypotheses have been proposed about the molecular mechanisms responsible for clustering genes. Duplication and divergence are at the basis of the Natal model, with metabolic pathways evolving in a step-wise fashion; the same idea is also central in the *Piece-wise* model where operons are assembled by clustering of sub-operons originated by in-tandem duplications of ancestral genes (Fani *et al*., 2005; Horowitz, 1945). In the *SNAP* hypothesis gene order rearrangement could be obtained by duplication events followed by selection during niche adaptation where non-selected genes are lost or inactivated (Brandis and Hughes, 2020). Operon formation could also take place on plasmids which may work as scribbling pads (Norris and Merieau, 2013). A recent proposal is based on consecutive reactions of IS insertion, deletion and excision, the IDE model (Kanai *et al*., 2022). Works on this topic often focus on the force driving gene clustering *or* on the way genes can get clustered. The former assume that molecular mechanisms for rearranging gene order exist and focus on plausible fitness functions, while the latter often assume some fitness function and focus on the ability of different types of genome rearrangements to build gene clusters (Ballouz *et al*., 2010; Castro and Brown, 2023). The lack of a single theoretical framework exhaustively accounting for the whys and hows of gene cluster assembly and maintenance suggests that different forces might be involved in this process and each, under appropriate conditions, may lead to gene clustering and operon formation, as also concluded by a real study-case (Martin and McInerney, 2009).

Here, we identify an additional drive for the *assembly* of gene clusters and operons, which is naturally active with varying intensity in all prokaryotic species. More specifically, we argue that metabolic gene clusters are evolutionary solutions that minimize the perturbations on metabolite homeostasis introduced by DNA replication and we show that gene clusters spontaneously form when we implement structural rearrangements explicitly, under this scenario.

Bacterial species have a more or less strict coupling of DNA replication with cell division. Species like *Caulobacter crescentus* and *Staphylococcus aureus* lie at one extreme, because they implement genetic programs that determine cell division just after the chromosome has been copied (Collier, 2012; Pinho *et al*., 2013), and are consequently strictly monoploid. These species can contain one or two chromosomes at most, and one replication fork. Other species, like for instance the two model organisms *Escherichia coli* and *Bacillus subtilis*, are able to switch from monoploidy at slow growth rates to mero-oligoploidy during rapid growth. In mero-oligoploids, several replication forks run concomitantly on the chromosome, enabling the bacterium to sustain division times shorter than the time for replicating the chromosome (Cooper and Helmstetter, 1968; Fossum *et al*., 2007). *E. coli* can for instance have up to 10 replication forks (Fig. 1a and b) (Pecoraro *et al*., 2011). The number of active replication forks can therefore change significantly in many species, but while an increased number of replication forks have been related to shorter division times in certain organisms (Trojanowski *et al*., 2018), the relationship seems to fail significance when distant taxonomic groups are analyzed together (Long *et al*., 2021). Other bacterial species can be oligo- or poly-ploids, owning from a few to tens or hundreds of full chromosomes, like many Cyanobacteria (Griese *et al*., 2011). In this case, depending on the species, replication can involve only one chromosome at a time, or many, and all of them are usually transcribed (Ohbayashi *et al*., 2019). Endosymbionts or giant Bacteria, show the most extreme ploidies as they can harbor up to hundred thousand chromosome copies (Nakabachi and Moran, 2022; Soppa, 2017). This issue can be investigated by using genome sequencing data, and calculating the ratio of the coverage at the *Ori* and *Ter* loci (*n*_*ori*_*/n*_*ter*_ ratio) (Fig.1). The artificial conditions used in the lab may be forcing the bacteria to growth rates that are rarely attained in the environment; yet, these ratios inform us on the potential ability to implement a mero-oligoploid mode of growth. The same quantity, often called the peak-to-trough (PTR) ratio, has recently been used to show that the growth rate of bacteria in the environment can change significantly (Joseph *et al*., 2022; Long *et al*., 2021), therefore cells experience a range of division times and must adapt their physiology to the ensuing challenges; approaches not based on this ratio, have confirmed this view (Haugan *et al*., 2018; Weissman *et al*., 2021).

**FIG. 1.**
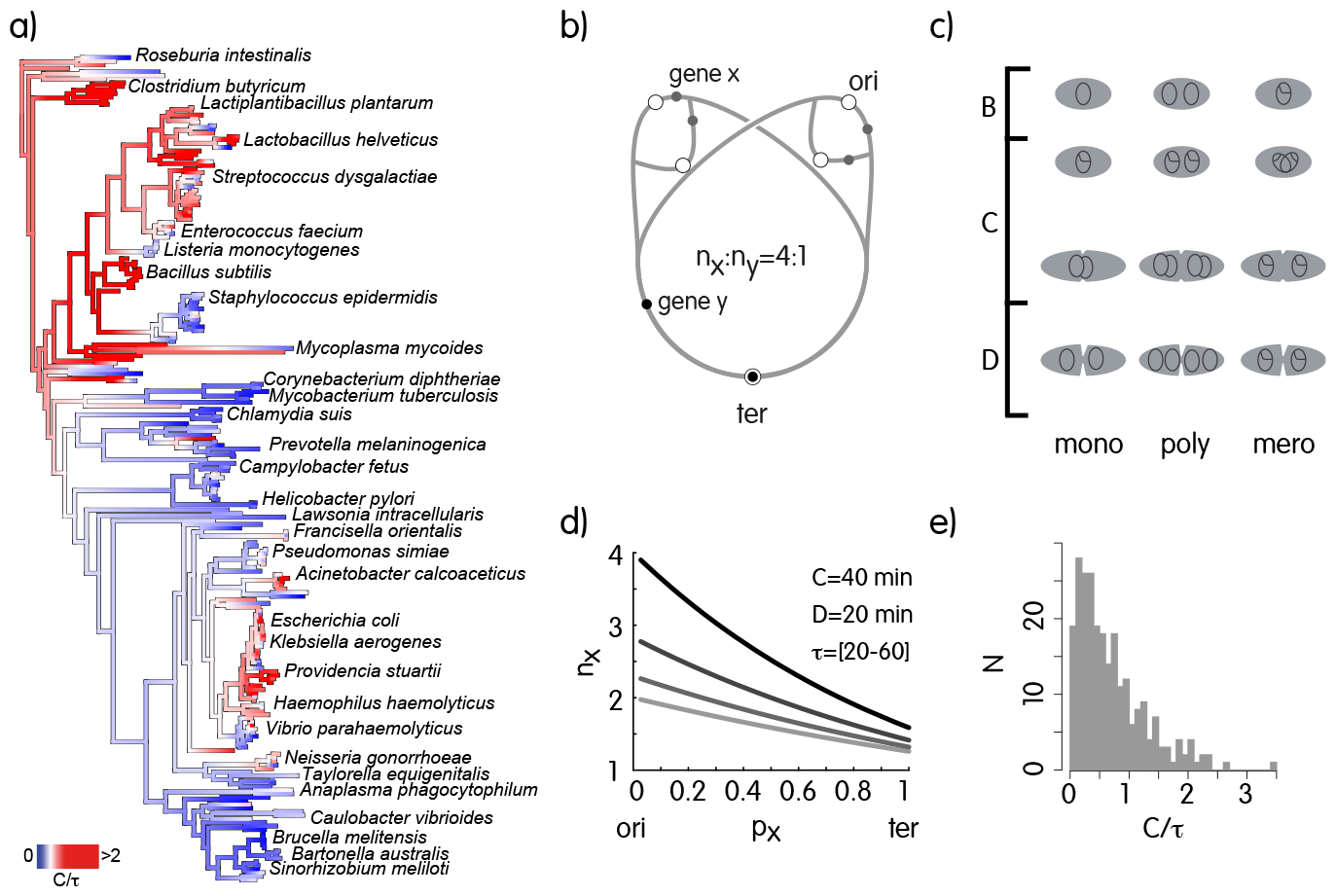
a) Mapping *log*_2_*n*_*ori*_*/n*_*ter*_= *log*_2_*PTR* = *C/τ* on a phylogenetic tree of Bacteria. Only a few species are indicated for clarity and *log*_2_ *n*_*ori*_*/n*_*ter*_ is truncated at 2 to highlight differences, whereas (*PTR*_*max*_ *≈* 10). Mono- and polyploid species have *PTR ≤* 2, with the maximum corresponding to situations where all cells are replicating all their chromosomes with a single active replication fork per cell. On the converse, mero-oligoploids can have ratios larger than 2, but not so at low growth rates and are therefore able to switch from one growth *mode* to another. Since these calculations exploits genome sequencing data, we assume that bacteria were grown in optimal conditions, therefore the observed ratios should correspond to maxima; this enables to detect species that are likely mero-oligoploid (in red). b) Replication and its effect on copy number of loci: With three replication forks as in the scheme, *x* and *y*, are in a 4 : 1 ratio but this changes with the number of forks; an *E. coli* cell can for instance be born with a single chromosome and after some time of exponential growth in a rich environment its progeny can have several genome equivalents and/or active replication forks. c) Different modes of growth in prokaryotes. d) Average copy number of loci as a function of the position on the Ori-Ter axis and division time (dark, shorter division time); e) Distribution of the *C/τ* ratios average estimates for species in the tree.

Recently, the use of single cell transcriptomic data demonstrated that replication imparts a clear effect on gene expression in *E. coli* and *Staphylococcus aureus*. Moreover, the same authors showed that this is also the case for *C. crescentus* gene expression data, in a synchronized population (Pountain *et al*., 2022). We therefore conclude that (i) loci experience changes in copy number depending on their distance from the origin during replication when the growth rate changes, and (ii) this affects the abundance of transcripts (Couturier and Rocha, 2006; Jaruszewicz-B-lonńska and Lipniacki, 2017), with maybe a partial buffering by global gene expression regulation, like super-coiling (Dorman, 2019).

Here, we provide theoretical considerations about the effect of active DNA replication on metabolic homeostasis. Under this view, replication can provide a selective force for driving functionally related genes at nearby loci in evolution, which we interpret as a fundamental pre-requisite for operon formation.

## RESULTS

### Metabolic consequences of chromosome replication

In this section we provide the theoretical foundations of our hypothesis by linking together chromosome replication and its possible effects on cellular homeostasis. We do this by integrating, for the first time, a widely adopted metric related to the average number of replication forks in cells from a population (the so called Peak-to-Trough ratio (Long *et al*., 2021) or simply PTR) with Metabolic Control Analysis (MCA), an established theoretical framework focused on understanding the control of metabolic fluxes.

If the PTR was constant (Fig.1b), the copy number of two genes *x* and *y* located at distant loci on the genome would also be constant. However, since the number of replication forks changes in time, those genes not only experience different multiplicities in time but also varying relative abundances. For instance, in Fig.1b with three replication forks, *x* and *y* are in a 4 : 1 ratio, but with one replication fork, the ratio becomes 2 : 1. Since the multiplicity of genes affects the abundance of transcripts (Pountain *et al*., 2022), the relative expression level of enzymes, especially when they are encoded by distant loci, is also a function of the number of replication forks. The formalization of the metabolic consequences of changes in the abundance of enzymes belonging to the same metabolic pathway is one of the fundamental results of MCA (Eq. 1) (Small and Kacser, 1993a):

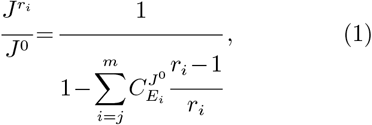

where the *J*s are steady state fluxes for the reference (^0^) or the new steady state, induced by changing the abundance of *m−j* enzymes in a pathway by different factors *r*_*i*_. 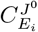is the *flux control coefficient* (FCC) (Kacser and Burns, 1973) of enzyme *E*_*i*_ on the pathway, defined in Eq 2.

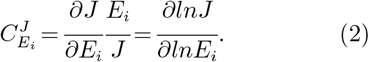

In practice, the FCC of enzyme *i* tells us about the fractional change in pathway flux elicited by a fractional change in enzyme abundance. FCCs are *systemic* quantities that can be measured only in the intact system. Nevertheless, the so-called *summation theorem* (Kacser and Burns, 1973):

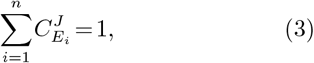

with *n* representing the number of enzymes of the pathway, constrains the range of values, at the same time introducing the fundamental concept that fluxes are not usually modulated by key *bottleneck* or *rate limiting* enzymes; instead, control is shared by many enzymes (Kacser and Burns, 1973). A consequence of Eq. 3, confirmed by *in vivo* measurements since early times, is that the flux control coefficient of an enzyme with respect to a pathway is small on average (Kacser and Burns, 1981; Small and Kacser, 1993a, b). Eq. 1 therefore shows that when all (i.e. the Summation theorem holds) enzymes in a pathway are changed by the same factor *r*_*i*_ = *r* the system relaxes to a new steady state where the flux is scaled by *r* and metabolite concentrations stay constant (perfect homeostasis). When the *r*_*i*_s are different, the ensuing change in flux instead depends both on the the FCCs and the *r*_*i*_s. In this case however, enzyme rates are not scaled proportionally, and therefore metabolite pools can also move to a new steady state. This effect on metabolites can be quantified by a relationship Eq. 1 that focuses on the deviation of metabolite concentrations when the abundance of one enzyme in a pathway is scaled by *r* (Small and Kacser, 1993b):

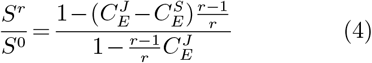

where 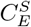 is the *concentration control coefficient* (CCC) of the enzyme over metabolite *S*, defined similarly to the FCCs (Kacser and Burns, 1973) and:

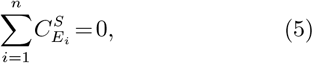

is the summation theorem for the CCC (Heinrich and Rapoport, 1974).We point out that when 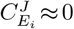 (i.e. the enzyme has negligible control on the flux) Eq. 4 gives (Fell, 2005):

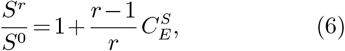

which shows - together with the fact that Eq. 5 does not limit the absolute value of the coefficients - that the manipulation of the abundance of an enzyme with negligible control over the flux of a pathway, can nonetheless perturb metabolite pools by an arbitrarily large factor (Fell, 2005).

Even with some limitations imposed by the FCCs (therefore by the system), cells manage to manipulate fluxes as a function of the demand while keeping metabolite concentrations within acceptable limits. This can theoretically be achieved by putting specific sets of genes under the control of a same regulator - as for yeast’s amino acid biosynthetic genes, that are controlled by *GCN4* (Albrecht *et al*., 1998). Alternatively, a similar achievement results from organizing genes in an operon. However, Prokaryotes have one major difference with respect to most Eukaryotes, and we are going to show that for this reason, the former strategy could be harder to accomplish. To better illustrate this, we integrate the change in multiplicity of genomic loci as a consequence of replication and the position of genes along the Ori-Ter axis into Eq. 1. This can be done by exploiting classical models that precisely connect the division time and the number of active replication forks (Cooper and Helmstetter, 1968; Helmstetter, 1996; Helmstetter and Pierucci, 1976). Indeed, when the duplication time (*τ* below, e.g. in hours), the time required to replicate DNA and the delay of cell division after the genome has been replicated (*C* and *D* in e.g. hours) are known, the number of copies of a locus *x* can be predicted by:

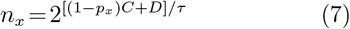

where *p*_*x*_ is the fractional position of the gene such that *p*_*ori*_ = 0 and *p*_*ter*_ = 1 (Helmstetter, 1996) (Fig. 1c). It is therefore easy to demonstrate that

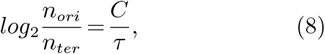

which relates a quantity that can be calculated from genome sequencing data to changes in division times, for instance when a bacterium shifts carbon source or in feast and famine cycles (Morin *et al*., 2020). We note that only mero-oligoploid species can have *n*_*ori*_*/n*_*ter*_ *>* 2 or *C/τ >* 1, i.e. a situation characterized by a DNA replication phase (C) longer than the division time.

A simple model for the abundance of a certain transcript *x* under the regulation of transcription factor *y* could be integrated with multiplicity of the *x* locus as:

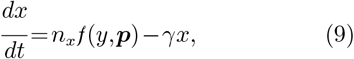

where ***p*** indicates the parameters of the regulatory function (*f*(*y*,***p***)), often having a type II or III functional response (e.g. Hill functions with *n* = 1 and *n ≥* 2, respectively).

The general solution for the steady state abundance of *x* is:

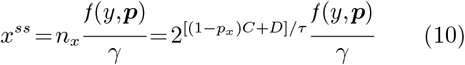

Which shows that the steady state level of a transcript is also determined by changes in the division time that should be common in the environment. By integrating this solution in Eq.1, we obtain:

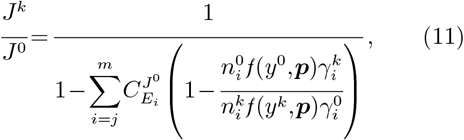

As expected, the amplification factor depends on both relative changes in (i) copy numbers of the loci, which are functions of division time, (ii) the activity of the regulator, and (iii) the degradation rate.

If all enzymes of the pathway are changed:

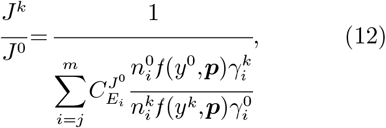

and if promoter characteristics and degradation rates are similar for all genes:

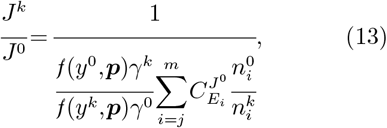

Finally, assuming the shift from condition 0 to *k* brings no regulatory and degradation changes at the same time not affecting *C* and *D*:

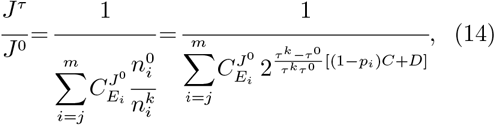

Eq. 14 highlights that, if a bacterium faces changes in division time across conditions, then the ensuing variation in copy number can perturb metabolic fluxes. As this situation may appear unreal, we highlight that even without such strong assumptions (Eq.13), replication still represents a confounding factor in the input/output response of classical transcriptional regulation. It also backs up criticisms to the idea that the optimization of promoter sequences of scattered genes is as good an option for Bacteria as for Eukaryotes: even with identical promoters, variations in *τ* can change the relative abundance of loci, hampering the evolutionary optimization of promoters, unless they are clustered on the genome. Genome rearrangements are an additional factor contributing to this difficulty, since by shuffling genes, they create novel configurations that may drastically decrease the optimality of promoters.

Since (i) Eq. 6 shows that even enzymes with negligible flux control coefficient can perturb metabolite pools by arbitrarily large factors, and that (ii) significant variations in metabolite concentrations in the cell can break down cellular homeostasis, we suggest that a possible solution worked out by evolution could be the formation of gene clusters and operons. In such a case, the *p*_*i*_s of all genes with control over a certain pathway flux (such that Eq. 3 holds) are very similar and Eq. 14 becomes:

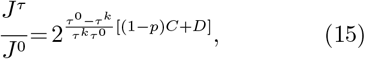

which is exactly the factor used to change all the enzymes, the one ensuring that metabolite pools don’t diverge from their previous steady state.

### Gene proximity minimizes variations in metabolite homeostasis

Let us now introduce a toy pathway to discuss more thoroughly the points raised above:

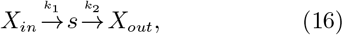

where *X*_*in*_ and *X*_*out*_ are external metabolites, and *k*_1_, *k*_2_ are the rates of two enzymes (with concentration *E*_1_ and *E*_2_) coded for by genes located at distances *p*_1_ and *p*_2_ from the origin. Using mass action we write this simple system as:

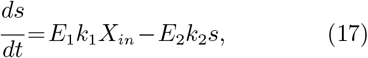

which can be solved analytically at the steady state (when *ds/dt* = 0):

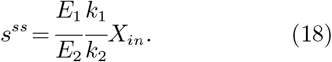

Enzyme abundances (*E*_*i*_s) can be replaced by the relation introduced in Eq. 7 and by derivation we can calculate the scaled sensitivity of the concentration of metabolite *s* with respect to changes in division time i.e. to variations in the average number of active replication forks per cell:

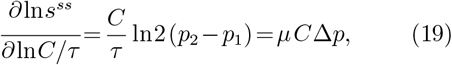

since by definition the growth rate is 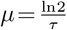.

The scaled sensitivity of metabolite concentrations with respect to the division time therefore depends on the difference in position of the two genes (Δ*p*) and the change in *C/τ*. Equivalently, the sensitivity depends on the relative delay in copying one gene with respect to the other 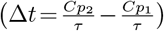. If *p*_1_ *>p*_2_, an increase in *C/τ*, or more intuitively a decrease in division time causes a depletion of *s* at steady state, while the opposite happens if the division time increases. However, if *p*_1_ *≈ p*_2_ the sensitivity tends to zero, making the pool independent from changes in division time.

This example provides additional theoretical basis for an effect on homeostasis introduced by changes in division time, whose degree depends on how genes are organized in the genome. It is therefore plausible that events leading to the minimization of those perturbations have a positive impact on the fitness of a cell and therefore increase its probability of fixation in the population. One of those mechanisms, as the above considerations suggest, is to group functionally related genes in a compact locus. To underline why, we refer to our toy model, and by scanning many positions for our two genes in a virtual 2 Mbp linear chromosome, we calculate a measure of the resulting variation in the abundance of metabolite *s* across a range of division times. Fig. 2a therefore confirms our theoretical prediction that, when the division time changes over a certain interval, metabolite homeostasis is maintained much better if the genes are close on the genome; it also shows that the magnitude of perturbation increases when genes are farther, suggesting a mechanism for evolution to select for intermediate steps in the construction of operons.

**FIG. 2.**
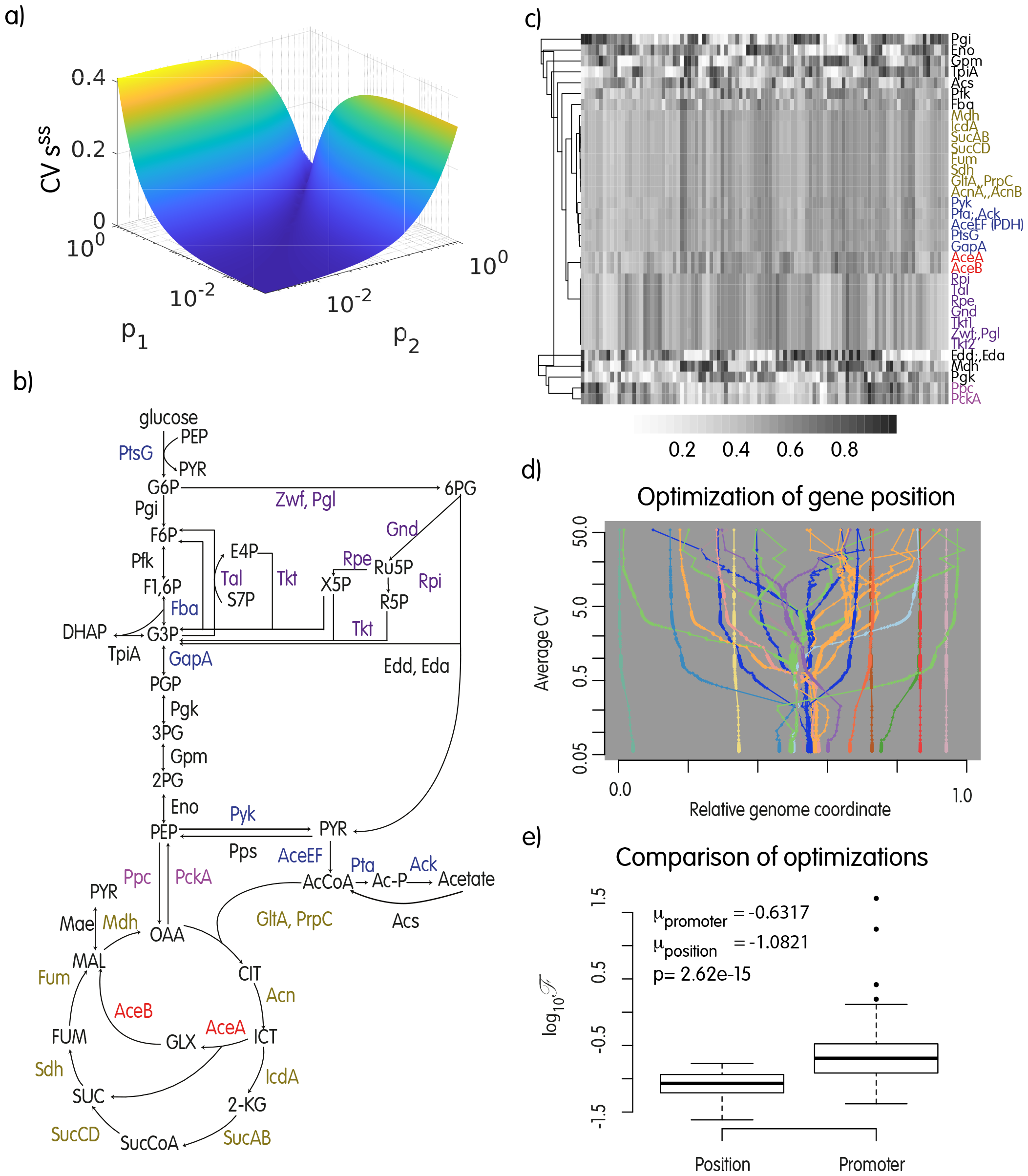
a) Scanning of the genomic positions for the two genes in the toy model (*p*_1_ and *p*_2_, expressed as a distance from the terminus). The z-axis is the coefficient of variation (standard deviation / mean) of metabolite *s* at steady state for 5 values of *τ ∈* [20,60]. When the two genes are close on the genome (along the diagonal in the plot), perturbations to the homeostasis of *s* consequent to the changes in multiplicity of the genes induced by the change in *τ* are efficiently buffered. b) The carbon utilization metabolic network implemented in the linlog model; c) The results of 100 independent optimizations as described in the text and started from random gene positions. Rows are the enzymes in the model; the colormap indicates the position of the corresponding gene at convergence i.e., same color in a column means the two genes are close on the chromosome. Groups of genes form similar clusters in different runs. d) All iterations of one optimization run from (c). Genes ending in the same genomic region have the same color. Sub-clusters can form and destroy during the optimization, but once the final configuration is found, it is stable in time. e) Comparison of the objective function at convergence for the optimization of gene positions and transcription rate constants (averages over 100 runs).

### Evolving operons *in silico*

The study of a very simple two-genes system provided support to the idea that in species where the number of active replication forks is partially decoupled from cell division, the operon might be instrumental for metabolic homeostasis.

To further test our hypothesis we introduce a more realistic metabolic model that we previously developed (Berthoumieux *et al*., 2011), comprising reactions collectively annotated as *carbon metabolism* in *E. coli* i.e. Glycolysis, Pentose-phosphate pathway, Krebs cycle, Glyoxylate shunt and Acetate excretion/import. The model (Fig. 2c) has 34 enzymatic reactions, 26 variable metabolites and two external metabolites (glucose and acetate). It is encoded as a linear approximation called *linlog*, that was introduced by (Visser and Heijnen, 2003; Visser *et al*., 2004) as an alternative to classical *Michaelis-Menten*-like rate functions; the latter are non linear, complicating parameter identification from the data; additionally, systems of even a few non-linear equations are usually not solvable analytically. In the *linlog*, each rate is modeled as a linear combination of logarithms of normalized metabolite concentrations, providing an analytical solution for the steady state (Eq. 21); the original parameters corresponds to the elasticity coefficients from MCA. Normalization of metabolites and fluxes is performed by their reference value in the original formulation, but here we include the reference level into the parameters to work on absolute quantities.

Using this approximation, rates are modeled as:

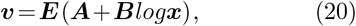

where ***E***_*r×r*_ is a diagonal matrix of enzyme abundances (modeled as in Eq. 7), ***A***_*r×*1_ and ***B***_*r×m*_ contain parameters derived from elasticities and reference steady state, and ***x***_*m×*1_ is a vector of metabolite concentrations. Given the stoichiometry matrix ***N***_*m×r*_, we can solve analytically the system for the steady state:

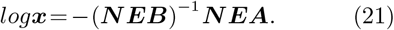

The model is not parameterized with respect to experimental data, but it has biologically meaningful parameters and therefore we used it here as a benchmark for testing our hypothesis in a realistic model. This is instrumental if one wants to test the systemic response to perturbations, while focusing on an abstract pathway removes the effect of the surrounding metabolic network.

Since matrix ***E*** contains enzyme abundances, we can simulate how the position of genes evolves in time when maintenance of metabolic homeostasis is the objective. Deviations from homeostasis can be summarized by the variation in metabolite concentrations when the division time changes. In brief, our experimental setting comprises variable division time, which can affect enzyme levels through the multiplicity of the corresponding locus on the genome, a chromosome with genes encoding the enzymes, and the metabolic model. We let gene position on the chromosome evolve to minimize perturbations to metabolite concentrations when the division time varies in *b* intervals. To this end, we use an optimization tool in R (nlminb) with the objective function shown in Eq. 22:

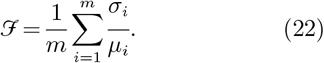

Where *µ*_*i*_ and *σ*_*i*_ are the mean and standard deviation of the concentration of metabolite *i* across the tested division times, and *ℱ* is therefore the average coefficient of variation of metabolite concentrations in the system when *τ* is changed. We stress here for clarity, that this simulation has no mechanistic basis for gene order reorganization and only considers the effect of genome structural variations on metabolic homeostasis by evaluating the objective function; it is therefore not informative on *how* genes actually move, but can provide information on the result in evolutionary time without the need for additional parameters. In paragraph *Simulation of structural rearrangements* we will show that a more realistic model gives a congruent answer but requires much longer simulation times.

In Fig. 2b, we show the outcome of 100 optimization runs. Several genes consistently form compact clusters at convergence (same shade of grey within a column) as expected by our theoretical considerations. Clusters formed at convergence of different optimizations are not always the same, with some of the genes ending in different or no cluster in different simulations, suggesting the existence of alternative, similarly optimal solutions, and/or an effect from the starting conditions of some genes. Globally, we observe a strong correlation of clusters formed in the optimizations and the pathway of the genes (compare with Fig.2c). Fig. 2d shows all the iterations for one optimization run, starting at the top with a random gene arrangement that evolves reaching a final configuration at the bottom line.

One alternative to the perturbations introduced by a varying number of replication forks concerns promoter evolution, therefore we implemented a similar optimization where genes have fixed coordinates, division time can change as in the above experiment, and enzyme levels are determined by a parameter under optimization to mimic the evolution of promoter characteristics. More precisely, enzyme levels are calculated as *E*_*x*_ = *n*_*x*_*k*, with *n*_*x*_ defined by the static position of gene *x*, the division times, and the *k* under optimization. Each iteration starts with random gene orders. Fig. 2e shows that promoters cannot provide the same degree of homeostasis as the optimization of gene positions. We conclude that the formation of gene clusters and operons might indeed represent an optimal evolutionary solution to cope with these metabolic perturbations.

The above considerations would suggest that the selective pressure driving the formation of compact gene clusters and operons is a function of the range of division times attainable by a certain organism; species may therefore experience different levels of selection towards the formation of operons, depending on the ability to change division time over a more or less wide range.

### Simulation of structural rearrangements

The optimization of gene position discussed above suggests that, during evolution, structural rearrangements of genomes should be able to build gene clusters, under the selective pressure discussed so far. However, the optimization algorithm used above does not explicitly account for specific molecular rearrangement and may therefore provide biased results. One major problem in a realistic scenario is for instance that rearrangements bringing some of the genes at nearby positions could split previously formed clusters, making the optimization of gene positions harder or even impossible. Therefore, we implemented a more realistic scenario where inversions and translocations are explicitly considered and we integrated it with our metabolic network model. Previous works studying the formation of clusters from a structural point of view often focus on a single mechanism (inversions or translocations) and on artificial definitions of pathways or other functional interactions of genes. Ballouz *et al*. (2010), have for instance demonstrated the ability of inversions to form gene clusters when the fitness is a function of the distance of two target genes belonging to the same virtual pathway (Ballouz *et al*., 2010). Their fitness function can be seen as an abstraction of what we formalized here, but the system is much simpler as the *pathway* is of two genes only, it is not embedded in a larger metabolic network, and the other genes are all neutral with respect to rearrangements. Therefore, we simulated a much more realistic scenario, with structural rearrangements taking place in a genome with 600 loci, comprising the 34 genes encoding the enzymes of the metabolic system. In summary, the simulation starts with a population of 10^5^ identical genomes; inversions and translocations take place with a certain constant rate. Once a structural rearrangement takes place in a genome, the fitness is calculated by exploiting the metabolic network at different growth rates, and accounting for the multiplicity of the gene calculated on the basis of its position with respect to the ori/ter axis like before. Once all genomes that underwent a structural rearrangement are processed, variants with a fitness less than the 50th percentile are removed from the population and the surviving genomes are sampled proportionally to their fitness. The new population then undergoes another generation with rearrangements, selection and so on. It is important to notice that we don’t define pathways in advance, therefore there is not a pre-defined gene order configuration to be achieved; instead, structural variants with higher fitness will increase in the population, and we follow the formation and number of gene clusters comprising target genes. We also stress that we did not reduce the probability of splitting previously existing clusters as done, for instance, by (Castro and Brown, 2023). During our simulations, each edge has therefore the same probability of being broken, also because additional simulations suggested that forcing the maintenance of clusters in our system, reduced the optimality of solutions, by forbidding better configurations (data not shown). The inclusion of the aforementioned mechanisms in our simulation framework did not affect the main outcome of our *in silico* evolution experiment. Indeed, as shown in Fig. 3, in all cases tested so far, one or more *pure* and/or *mixed* clusters (as defined in *Methods*) always formed, thus confirming and extending our hypothesis. It should be noted that the degree of clustering is indeed smaller in the more realistic case, but we demonstrated that the selective force proposed in this work is able to select for gene configurations - also comprising a certain number of *clusters*, that enable a huge improvement in homeostasis.

**FIG. 3.**
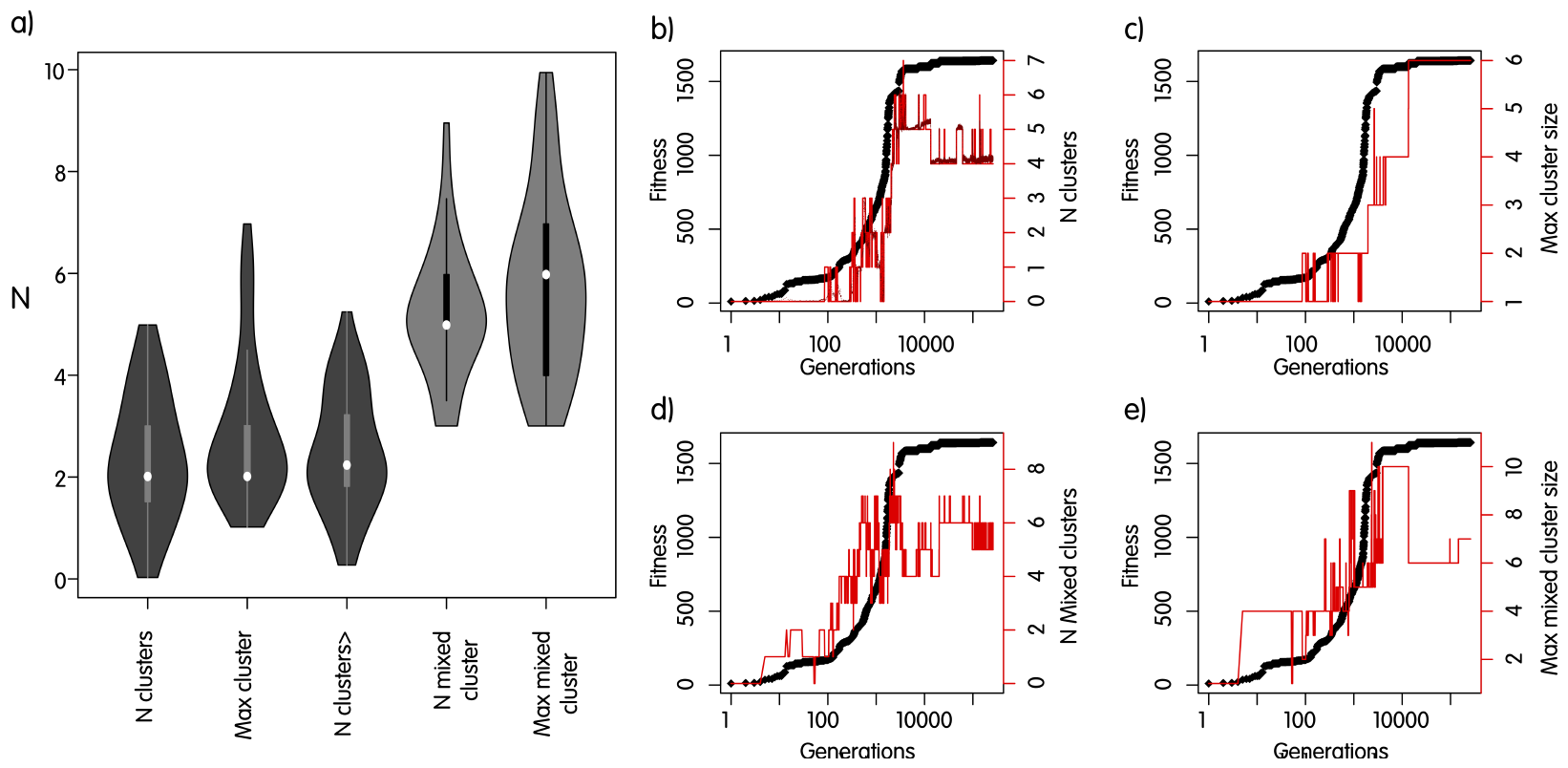
Simulations of gene order evolution with inversions and translocations. a) Counts and lengths of clusters in 50 independent simulations ran for 100k generations; b-e) Results for one specific simulation after 250k generations. In black, the fitness of the fittest strain in the population. b) Number of clusters containing only metabolic genes in the fittest strain. A cluster is defined as a contiguous stretch of genes. Dark red dots, mean number of clusters in the population. c) Number of genes in the largest cluster. d), e) as b) and c) but for mixed clusters (see definition in *Methods*).

### Degree of compaction of functionally related genes correlates to the PTR

One testable consequence of our hypothesis is that species able to achieve larger PTR should experience a more effective selective pressure for clustering functionally related genes. As all species can attain long division times, we can consider the average PTR in laboratory conditions as a proxy of the division time range a species can potentially experience. Two confounding factors in this attempt are horizontal gene transfer and the evolutionary rate of the PTR itself: metabolic operons are easily transferable self-contained modules that are likely advantageous even in species with a very low average PTR; additionally, the PTR calculated from genome sequencing reads is a snapshot of present-day organisms, while the degree of gene clustering in genomes reflects a much longer and unknown evolutionary path. Nonetheless, the fact that bacterial species in optimized conditions attain widely different growth rates represents a genuine *ability* : when replication of the chromosome and cell division are genetically entangled, like in *Caulobacter* there’s a natural bound to the division time since *τ ≥ C* holds. On the converse, the shortest division time must be defined otherwise in species where the coupling of DNA replication and cell division is weaker.

With these drawbacks in mind, we tested our hypothesis by deriving a *Proximity Score* (PS) that summarizes the degree of compaction of functionally related metabolic genes in complete genomes (see the complete list at https://github.com/ComparativeSystemsBiologyGroup/OperonEvolution). For each organism and functional category in KEGG we retrieved the corresponding genes, sorted them by position and recorded the distance separating consecutive ones. After processing of all pathways we calculated the 25th percentile of such distances; our PS is the logarithm base 2 of the inverse of this number, such that larger values correspond to more compact configurations. The PS was then compared to the *C/τ* ratio obtained from genome sequencing data, by using both regression models and t.tests in R.

Fig. 4a shows the existence of possible covariation of the two traits across the phylogenetic tree in a qualitative way; Fig. 4b confirms the presence of a significant relationship of the PS with *C/τ*, when using linear regression models; the coefficients are significant when considering all organisms, or the Proteobacteria, but not the Firmicutes alone. When significant, the model explains around 10% of the total variance in the data, which is a consistent fraction if we think about the many additional forces that act on genome organization in much shorter evolutionary times. In Fig.4c we strengthen this idea by showing that species with *C/τ >* 1 with *p ≤* 0.01) also have a larger PS. Since the Firmicutes have PS and *C/τ* ratios that are significantly above those in Proteobacteria, we additionally show in Fig.4d that the difference is significant even if we limit the test to Proteobacteria. The situation of the Firmicutes may seem in contrast with our hypothesis. However, when comparing their PS and PTR to the Proteobacteria, we found they are both significantly larger (*p<* 2.2*×*10^*−*16^) for both PS and PTR (*p<* 3.8*×*10^*−*4^). This suggests that the Firmicutes likely have a tendency toward high PTR values since their common ancestor, which may explain the thorough optimization of gene organization in this group.

**FIG. 4.**
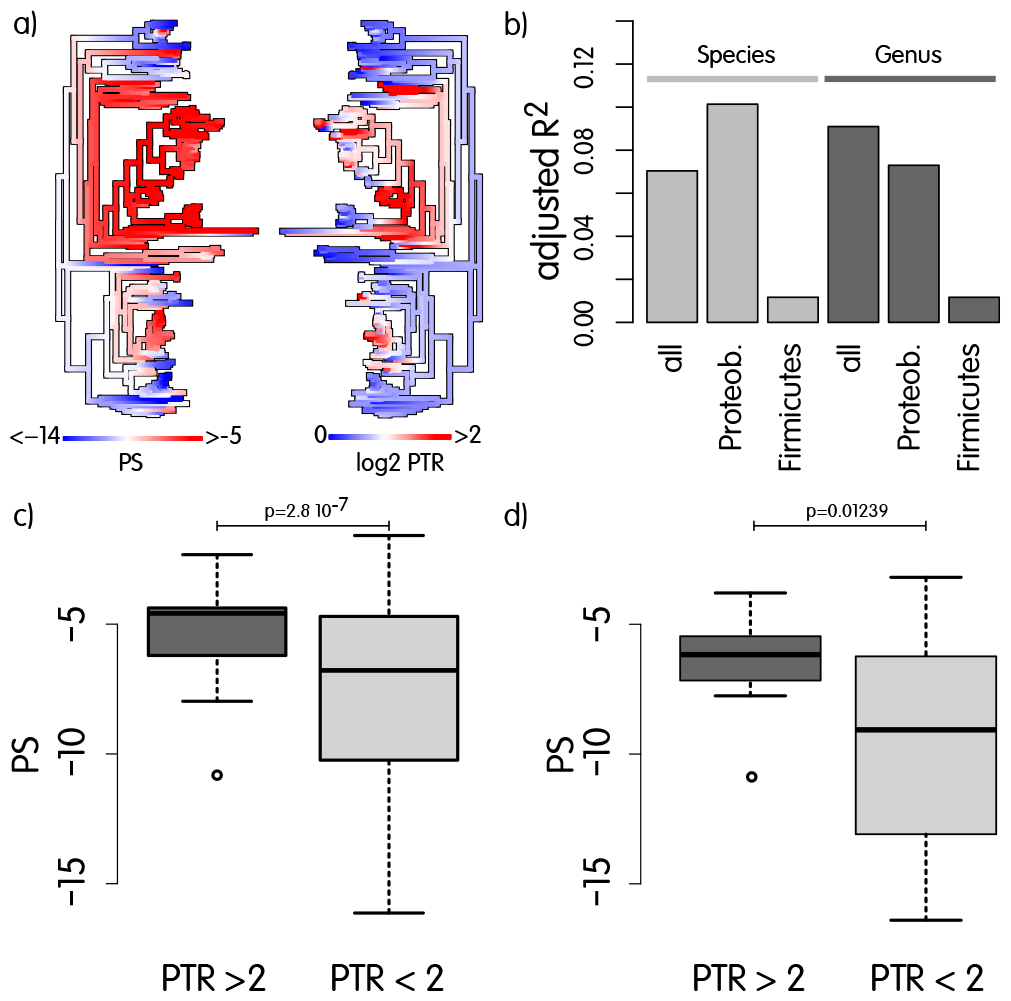
a) Face to face phylogenetic trees with mapped ancestral reconstructions of Proximity Score (PS) and *log*_2_*PTR*. b) Barplot of the adjusted *R*^2^ for linear regression models using PS as a predictor of *log*_2_*PTR*. All regression coefficients for PS are significant except for Firmicutes (data not shown). *Species* correspond to models exploiting all the organisms in the dataset and *Genus* means that models are build on averaged values across species belonging to the same genus. We also built models for Proteobacteria (*N* = 46) and Firmicutes (*N* = 29) as they cumulatively represent 86% of the species under analysis; c) boxplot showing that species having *p ≤* 0.01 when testing for *PTR ≥* 2 tend to have significantly larger PS. Firmicutes have significantly larger PS (*p<* 2.2*e −* 16) and PTR (*p* = 0.0003766) with respect to the Proteobacteria, and this may affect the above pvalue, but d) which is limited to Proteobacteria, shows that the test is still significant without Firmicutes.

## CONCLUSIONS

In this paper we propose and explore the novel hypothesis that metabolic gene clusters evolve to face metabolite pools perturbations introduced by DNA replication. Using a set of simulations of increasing complexity that combine MCA with a simple model of gene multiplicity during replication, we provide the theoretical basis for our hypothesis and suggest the existence of a selective pressure that likely contributed to gene clustering. In doing so, we also disclose a plausible mechanism through which other hypotheses can play a significant role, as they require genes in close proximity. The improvements of this work with respect to previous ones is that since flux and concentration control coefficients are systemic properties, we relied on a realistic metabolic system and we integrated it with a genome evolutionary model that explicitly takes into account structural rearrangements. Another important aspect of this work, is that we are not using an artificial definition of a metabolic pathway but, rather, we define a metabolic system potentially containing many overlapping *pathways*, and we let gene organization evolve on its own based on the effect on metabolite homeostasis. This complicates and slows down considerably the evolutionary process because many structural rearrangements that move some of the genes closer, likely split previously formed clusters. However, this is exactly what happens in real genome evolution where all genes somehow exert an effect on fitness and are therefore not completely free of moving; the genomes where an event breaks a pre-existing cluster will survive if this is compatible with their fitness. Even with these complications, we were able to observe the formation of clusters and a strong increase in the fitness consequent to genome rearrangements. Therefore our simulations have shown that introducing the PTR as a controller of enzyme abundances in a metabolic model and fixing an evolutionary meaningful objective such as metabolite homeostasis, led to the formation of gene clusters. One possible limitation of our hypothesis relies in the existence of a critical distance separating two genes (*d*_*crit*_) such that variants with the genes closer than this threshold do not get any appreciable fitness increase. In brief, when *d*_*a,b*_ = *d*_*crit*_ further reductions in distance become practically neutral. In this scenario, additional compaction may be indirectly selected in the population by the negative selection of variants where 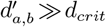 if this is associated to a significant decrease in fitness. Moreover, when there are more than two clustered genes, the critical threshold refers to the largest distance, and therefore further compaction within the cluster could still happen. Ultimately, we consider our hypothesis instrumental for driving genes at short distances, a prerequisite for many other hypothesis for operon formation. Indeed, once genes get close, other mechanisms can take over: production of functionally related proteins in a smaller volume may provide an advantage, and/or the cluster may become a single transcriptional unit. This would make the cluster more *resistant* in evolutionary time, such that the selection for specific transcription patterns could drive the removal of functionally unrelated genes (Fig. 5).

**FIG. 5.**
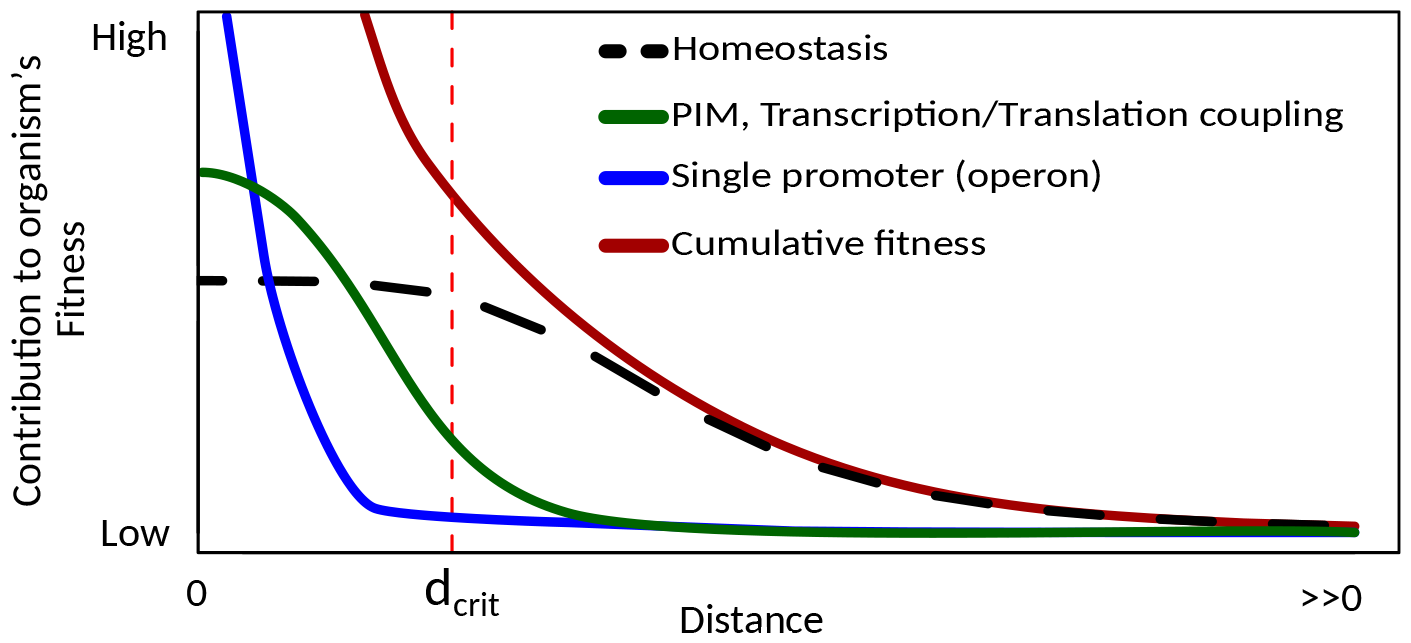
The selective force for better homeostasis is able to explain gene clustering, but once a certain critical threshold *d*_*crit*_ for inter-gene distance is reached, further compaction can be driven by other hypotheses as well. Profiles and relative contributions are arbitrary.

Our theoretical predictions suggest that the larger the average number of replication forks, the stronger the selective pressure toward gene clustering. By deriving a measure of the degree of compaction of functionally related genes for many species, we indeed confirmed that mero-oligoploid species have an organization of functionally related metabolic genes that is significantly more compact.

We believe that our hypothesis has several important repercussions on the way we conceive metabolic operon evolution on the one side and how we see ploidy and DNA replication in Bacteria on the other. Indeed, here we propose that the selective pressure toward gene clustering would be a function of the number of replication forks (summarized here by the PTR) such that mero-oligoploids could be considered as major *gene cluster formers*, while others, especially mono-ploids would be subject to reduced pressures in that sense. Nonetheless, fully formed operons still provide significant advantages to the latter, and horizontal gene transfer might be responsible for the diffusion of operons.

Not all cellular processes might benefit from the clustering of their genes into operons as a consequence to the unbalance described in this work; functionally related genes involved in other cellular processes than metabolism might behave differently but wherever two or more proteins work in precise ratios, such as in macro-molecular complexes, a similar effect could be at work since sub-units produced in excess would represent a waste of energy, which is in agreement with the results described by (Castro and Brown, 2023).

## METHODS

### PTR calculation, models and optimizations

Identification of origin and terminus for each genome was done with function *oriloc* from package seqinr (Charif and Lobry, 2007). Coverage around the two loci was obtained by first extracting a 50kb region centered on the *Ori* (*Ter*) locus, and then mapping publicly available genome sequencing data from SRA using Salmon (Patro *et al*., 2017) in mapping mode. Only species with at least 5 sequencing libraries available were considered. Coverage was not normalized because we focus on the ratio at the two loci. The R function t.test was used for evaluating *log*_2_*PTR* larger than 0 (*PTR>* 1) or larger than 1 (*PTR>* 2), the latter identifying mero-oligoploid species), and also provided 95% confidence intervals.

The optimization-based simulation was carried out using the R function *nlminb* and using as objective function the average coefficient of variation in metabolite concentrations when division times are changed, as explained in the main text. Genes can move on the genome under the hypothesis that genome organization should minimize perturbations to homeostasis, but no mechanistic model for genome organization evolution is explicitly implemented. The two gene-system was implemented in Matlab. The scripts implementing the models are available at https://github.com/ComparativeSystemsBiologyGroup/OperonEvolution.

### Simulation of gene order evolution

A genome is defined as a circular graph with 598 genes plus a vertex for the origin and one for the terminus. The origin is defined *a priori*, while the terminus is selected to be approximately equidistant from the origin. During the simulation, genomes where the minimum Ori-Ter distance is less than 75% of the original distance as a consequence of rearrangmenents, are assigned a very low fitness and are consequently removed from the population. All genes are separated by 10k nucleotides, to adjust the size of the artificial genomes to the Helmstetter and Cooper model. The simulation starts with 100k identical genomes; at each generation genomes can undergo structural rearrangements with a certain rate. In the simulations presented here, we used two rates for inversions, *p*_*inv*_ =[0.005,0.0025] and *p*_*transl*_ = *p*_*inv*_*/*2. Once an inversion occurs, it moves blocks of *N* (*mean* = 5,*var* = 9) genes, while translocations of *N* (*mean* = 3,*var* = 9). After all genomes undergoing an event are processed, fitness values are re-calculated, and all variants below the 50th percentile are removed.

The remaining are (i) compressed into unique structural variants and (ii) sampled proportionally to their fitness to re-generate the population of 100k individuals for the next generation. During the simulation, we keep trace of the number of *pure* clusters defined as contiguous stretches of at least 2 genes from the metabolic network and the maximum size of a cluster. We also define *mixed* clusters as clusters with metabolic genes at the extremities and at most two consecutive non metabolic genes inside.

### Gene clustering analysis and proximity score calculation

KEGG, the Kyoto Encyclopedia of Genes and Genomes (Ogata *et al*., 1999), provides a collection of pathway maps representing our knowledge of the molecular interaction, reaction and relation networks for a number or biologically relevant areas of research, such as Metabolism or Cellular Processes. Each KEGG pathway also contains manually defined functionally tight gene sets of different types and scopes. Metabolic modules are assigned to the *Pathway module* category, on which we focus our attention in this work. For every species under analysis (181), we obtained all gene-to-gene distances within each Pathway module and then calculated the first quartile after processing all modules. This number represents the gene-to-gene distance such that 25% of all distances are smaller. The logarithm base 2 of the reciprocal of this number is the Proximity Score (PS) of a species that we contrast to the PTR. Script implementing these analyses and the associated data are available at https://github.com/ComparativeSystemsBiologyGroup/OperonEvolution.

